# A computational solution to improve biomarker reproducibility during long-term projects

**DOI:** 10.1101/483800

**Authors:** Feng Feng, Morgan P. Thompson, Beena E. Thomas, Elizabeth R. Duffy, Jiyoun Kim, Shinichiro Kurosawa, Joseph Y. Tashjian, Yibing Wei, Chris Andry, D.J. Stearns-Kurosawa

## Abstract

Biomarkers are fundamental to basic and clinical research outcomes by reporting host responses and providing insight into disease pathophysiology. Measuring biomarkers with research-use ELISA kits is universal, yet lack of kit standardization and unexpected lot-to-lot variability presents analytic challenges for long-term projects. During an ongoing two-year project measuring plasma biomarkers in cancer patients, control concentrations for one biomarker (PF) decreased significantly after changes in ELISA kit lots. A comprehensive operations review pointed to standard curve shifts with the new kits, an analytic variable that jeopardized data already collected on hundreds of patient samples. After excluding other reasonable contributors to data variability, a computational solution was developed to provide a uniform platform for data analysis across multiple ELISA kit lots. The solution *(ELISAtools)* was developed within open-access R software in which variability between kits is treated as a batch effect. A defined best-fit Reference standard curve is modelled, a unique Shift factor “S” is calculated for every standard curve and data adjusted accordingly. The averaged S factors for PF ELISA kit lots #1-5 ranged from −0.086 to 0.735, and reduced control inter-assay variability from 62.4% to <9%, within quality control limits. S factors calculated for four other biomarkers provided a quantitative metric to monitor ELISAs over the 10 month study period for quality control purposes. Reproducible biomarker measurements are essential, particularly for long-term projects with valuable patient samples. Use of research-use ELISA kits is ubiquitous and judicious use of this computational solution maximizes biomarker reproducibility.

## Introduction

In virtually every research project with real or potential clinical application, biomarkers provide valuable data to monitor presence or progression of disease, as well as therapeutic susceptibility or efficacy. Biomarker data monitor defined outcomes and there is considerable discussion about whether investigators should disclose incidental research findings to study participants [1–3]. Intrinsic to this discussion is the need for reproducible study data and this presents challenges, particularly with long-term studies. Data rigor and reproducibility is a systemic problem [4] and a priority issue for the NIH. It is an analytic challenge for long-term studies because research laboratories often do not have standardized operations nor validated biomarker assay reagents that adhere to the quality assurance and quality control standards for diagnostic use as required by the Clinical Laboratory Improvement Act (CLIA; www.cdc.gov/clia).

Protein biomarkers are measured frequently in plasma, serum or other matrices by solid phase Enzyme-Linked Immunosorbent Assay (ELISA) methods in which the antigen of interest in the sample is bound by antibodies and the amount of bound antigen is proportional to the signal strength that develops in the assay. There have been only 136 *in vitro* diagnostic ELISA kits or kit components cleared or approved by the FDA since 2000, but there are hundreds of commercially available ELISA kits labeled for “research purposes only” from dozens of vendors. Typically these are vetted by the manufacturer for sensitivity, selectivity, intra/inter-assay variability, stability and storage needs, but they are not required to adhere to federal CLIA guidelines. Lot-to-lot variability between ELISA kits is either not relevant or is manageable for short-term projects, but challenges arise for quality assurance when multiple lots of research ELISA kits are used in long-term projects. A research laboratory may operate under NIH biomarker recommendations [5] and Good Clinical Laboratory Practice guidelines with appropriate training, auditing, assay validation and proficiency testing [6], but it does not have jurisdiction over kit reagents controlled by the manufacturer. In general practice, expected inter-assay coefficients of variation (CV) for ELISA standard curves will be within 10~20%, but if a commercial ELISA standard curve suddenly shifts significantly with a new kit lot and validated control data shifts outside limits, then the current patient biomarker concentrations cannot be compared with those quantified months earlier. An experienced laboratory can rescue data from one ELISA with a failed standard curve [7], but long-term quality assurance poses other challenges. Either all the samples have to be re-assayed, which introduces variables such as freeze/thaw issues, or the data discarded, all of which wastes precious patient samples and resources.

This was encountered by our research group with one commercial ELISA kit during an on-going two year project to evaluate effects of pre-analytic variables on plasma thrombosis biomarkers in patients. The project developed thirty-five standard operating procedures (SOPs) that define and document operations from blood acquisition to transport, processing, assay and storage. Nine biomarkers are quantified by research-use ELISA kits. For one biomarker, biomarker “PF”, after months of reproducible assays, we observed a significant shift of our standard curves and internal control results when assayed with new kits lots. The scope of the problem was revealed by review of data from 5 kit lots over 10 months and 65 ELISA plates. Rigorous review by the quality assurance team did not identify laboratory or operational pre-analytic contributions and similar changes were not observed with the other ELISAs. The project had encountered an unexpected analytic variable and the data from 420 plasma samples collected over 10 months could not be compared. The manufacturer was responsive but ultimately unable to resolve the problem.

To rescue our patient data, we developed a computational solution with a sufficiently generalized approach such that it may be used by others facing a similar situation. In the solution, the lot-to-lot variability in ELISA kits is treated as a batch effect, and a defined Reference standard curve is modelled with either a four-or five-parameter logistic function. Based on this Reference curve, a Shift factor (“S”) can be calculated and applied retrospectively to every standard curve from every ELISA plate over many months, and the biomarker concentrations for that plate are adjusted accordingly. In this way, the data collected from many ELISA plates over many months can be compared on a uniform platform. Once instituted, calculating the Shift factor for each ELISA standard curve or each kit lot provides an expedient way to rapidly monitor standard curves as a quality assurance metric and to facilitate data management.

## Materials and methods

### ELISA kits

Biomarker ELISAs with at least two kit lots were analyzed for the current study. ELISAs for human P-selectin/CD62P, human myeloperoxidase and human plasminogen activator inhibitor-1/serpin E1 were provided by R&D Systems (Minneapolis, MN, USA). The ELISA kit vendor for the biomarker of focus for the current study (biomarker “PF”) is not provided for discretionary reasons. Five lots of the biomarker PF ELISA kits were received over a 10 month time period. All ELISA kits were a standard 96-well format, sandwich antibody-based ELISA designated “for research purposes”.

### Assays and Equipment

ELISA kits were stored at 4 ± 2°C in a cooler (Helmer Scientific, Noblesville, IN, USA) equipped with alarmed wireless external temperature monitoring (SensoScientific, Inc., Simi Valley, CA). Temperature logs were reviewed and constant temperatures without drift were verified. All kits were used within the manufacturer’s expiration date. Plasma samples stored at −80°C were thawed just before assay in a 37°C water bath for <10 minutes, gently mixed, and kept on ice. Plasma samples were assayed according to a detailed standard operating procedure (SOP) for each biomarker ELISA that includes the manufacturer’s procedural steps. The SOP also included required documentation for every ELISA plate for operator, date, plate ID, critical reagents (date received, lot number, dilution/concentration, expiration date), incubation times (date, start/stop times, temperature), equipment (manufacturer, model, serial number) and a section to document any deviations from the SOP. Every plate included kit standards prepared according to the manufacturer’s instructions. The standards were added to triplicate wells in the first three columns of the provided 96 well plate (columns A-C, rows 1-8 using standard plate designations).

For every biomarker and every plate, an internal spiked plasma-based control sample (BMC Control, see below) was included in triplicate wells. Samples were added to the plates in with calibrated pipettes, washing steps were performed with an automated plate washer (Biotek model Elx50, Winooski, VT) and developed color was quantified by measuring optical density (O.D.) at the appropriate wavelength with a microplate reader (VERSAmax; Molecular Devices, Sunnyvale, CA). The acceptable coefficient of variation (CV) of triplicate wells for each standard, control or unknown was ≤15%. For biomarker PF, incubations with samples and detection antibody were done at 37°C in a dry incubator per the manufacturer’s instructions and perimeter wells were not used for patient samples to prevent possible evaporation complications due to the elevated incubation temperature. Other ELISAs were performed at room temperature. Initial data analyses were done with SoftMax Pro version 7 (Molecular Devices).

### BMC Control Preparation and Storage

Every ELISA plate included a human pooled plasma control sample that had been spiked with supplemental biomarker and designated the BMC Control. For each biomarker, the BMC Control was made in bulk volume according to the respective ELISA SOP and stored at −80°C in small aliquots for single use. For biomarker PF, lyophilized human pooled citrated plasma (Sigma Aldrich, catalogue P9523-5ML) was reconstituted with deionized water at room temperature for at least 15 minutes with gentle mixing, diluted with appropriate buffer to the same ratio as the patient samples and then spiked with reconstituted biomarker PF standard prepared from the same manufacturer’s kit but purchased expressly for this purpose. On the assay day, a BMC Control aliquot was thawed just before assay in a 37°C water bath for <10 minutes, mixed gently and added to each plate in triplicate wells. Two preparations of BMC Controls were made and aliquoted for storage: one in August 2017 (C1) and one in October 2017 (C2). Both preparations were made with PF standard from kit lot #1. The mean O.D. ± S.D. for C1 = 0.759 ± 0.095 (CV=13%, n=26 plates) and 0.672 ± 0.062 (CV=9%, n=16 plates) for C2.

### Data Analysis and Derivation of Shift Factor “S”

The approach was implemented in the statistical package R, an open software environment for statistical computing and graphics that accepts ELISA optical density data and standard concentration data for calculation of a best-fit Reference standard curve. The lot-to-lot variability is modelled as a fixed batch effect calculated as the difference between each plate’s standard curve and the Reference curve. This difference is designated the Shift factor “S”. An adjusted plate standard curve is derived using the S factor and used to adjust the biomarker concentrations.

The Reference and standard curves are fitted with four-or five-parameter logistic functions (4pl, 5pl). These functions are well established models to relate analyte concentrations to their response signal intensities in immunoassays [8–10]. The 5pl has the form of:

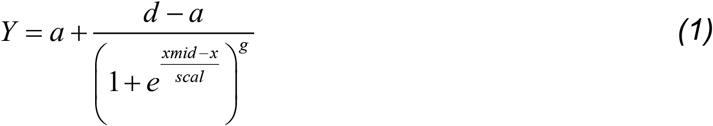

where *Y* is the signal intensity of measurement (OD in ELISA assays); *x* is the log-transformed concentration of analytes; *a* and *d* are the lower and upper asymptotes of signal intensity, respectively; *xmid* is the x value of the curve’s inflexion point; *scal* is the scale parameter or the inverse of the slope of the curve at the inflexion point (*x = xmid*); *g* is the factor controlling the curve asymmetry. When *g* takes a value of 1, the 5pl becomes the 4pl function.

The 5pl could also be written equivalently as a non-logarithm or exponential form,

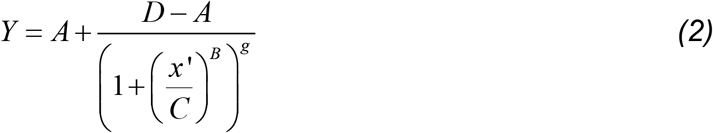

where *Y* and *g* are identical to the parameters in eq. (1); *x*’ is non-log transformed concentration and equal to *e^x^*; *A* and *D* are identical to its equivalent lower-case letter parameters in eq.(1); *B* and *C* are equivalent to *xmid* and *scal*, but have an exponential value of them. In this study, the log-form equation is used for implementation.

To analyze data from multiple plates or lots of ELISA kits, batch effects have to be modelled and corrected for data comparison. Many biological or technical factors could impact immunoassay reproducibility and lead to batch effects [11–15]. Lot-to-lot variability between PF ELISA kits is among such factors. We proposed to model and correct it as a fixed batch effect. It first assumes the lot-to-lot differences mainly result from the variable quantitation of standard analyte concentrations, which can expressed as:

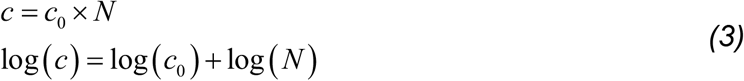

where *c*_0_ and *c* are the real and provided standard analyte concentrations from the manufacturer, respectively, and *N* is the fold difference between them. We can use *x, x*_0_ and *S* to replace log(c), log(c_0_), and log(*N*) and rewrite the above log-transformed equation as:

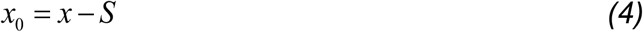

Therefore the statistical model of the standard curves can be written as:

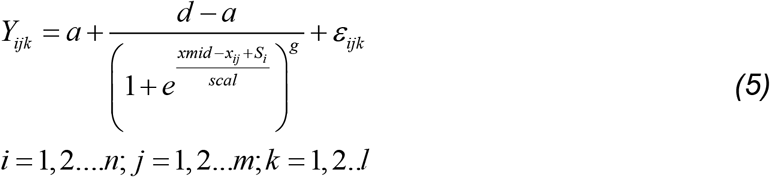

where *a, d, xmid*, *scal* and *g* are the parameters for the 5pl as in eq.(1); *Y_ijk_* is the observed signal intensity (measured OD in ELISA); *x*_*ij*_ is the log-transformed concentration of the *j^th^* standard analytes in the *i^th^* batch; *S_i_*, the Shifting factor, is the log fold difference between the known concentration and the true one; *i* and *n* are the *i^th^* and total batch number, respectively; *j* and *m* are the *j^th^* and total number of standards, respectively; *k* and *l* are the *k^th^* and total number of measurements; *ε_ijk_* is the random error for each measurements. This equation can be further rewritten into:

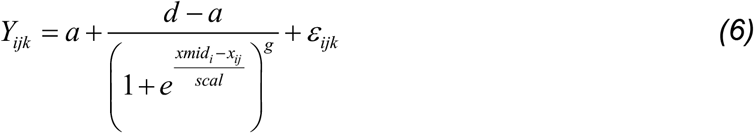

where *xmid_i_* = *xmid* + *S*_*i*_ and all other parameters are the same as in eq. (5). As a result, the model indicates that the standard curves of the same batch differ from each other as a result of random errors of measurements, while differences between curves from different batches is ascribed to inaccurate quantitation of analyte concentrations. Furthermore, the standard curves from different batches follow the 5pl functions (or 4pl) with the identical parameters of *a*, *d*, *scal* and *g*, but different *xmid*. The differences are defined by the Shift factor, *S_i_*, which is estimated through the non-linear regression together with other 5pl parameters.

To do the batch normalization/correction, the analyte concentrations in unknown samples are first estimated based on unadjusted standard curves and then the Shift factor *S* of the batch is applied to obtain the final quantities:

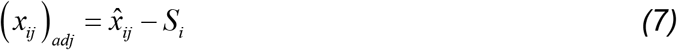

where *x* is the log-transformed analyte concentrations in the unknown sample, *i* and *j* are the batch and standard number as in eq.(5), respectively.

It might not be possible to know the precise concentrations of analyte in the standard samples (*c*_0_ in eq.(3)) without other validation methods, such as proteomics [16]. Therefore, we designated one batch as the Reference batch, in which the analyte concentrations of the standard samples were treated as accurate, and from that we estimated the Shift factor *S* of all other batches relative to it. The data from 28 standard curves from PF ELISA kit lot #1 (batch #1) was used to calculate a 4pl reference curve for biomarker PF. For other biomarkers, at least four representative standard curves from at least two lots and three operators were chosen to model the best-fit 4pl Reference curve.

## Statistical Analysis

Student’s t-tests were performed for analysis of inter-assay differences. Analysis of variance between groups was performed with Bonferroni post-test. P<0.05 was considered significant.

## Software Availability

An ELISA data analysis tool *(ELISAtools)* with the ability to correct batch effects has been implemented in the statistical R software (version 3.5.1) and is available freely for academic use at https://github.com/BULQI/ELISAtools. Instructions for calculation of S factors is provided in Supplemental Methods.

## Results

Procedurally, the ELISA manufacturer’s directions were followed for reagents, buffers, assay temperature, reagent incubations and wash times. Operationally, these instructions were supplemented with documentation for each plate that included operator, kit reagent lot numbers and clock times for reagent additions and incubation periods. Including a BMC control sample on each plate permitted comparison of data over months (Fig 1). Control optical density (O.D.) readings were consistent with time even with different ELISA kit lots (Fig 1A,B). Myeloperoxidase data (lots #1,2) is shown as a comparison with biomarker PF (lots #1-5). Observations were similar for the other seven biomarkers (data not shown). Myeloperoxidase O.D.s were slightly higher for lot #2, but calculated antigen concentrations were stable (Fig 1C). In contrast, calculated PF concentrations in the BMC controls (preparations C1 and C2) decreased by an average 62.4% between ELISA kit lot #1 and lot #5 over the time (Fig 1D), exceeding our quality control limits. This disconnect between O.D. readings and calculated PF biomarker concentrations over time raised problems for the 420 patient samples already analyzed.

**Fig 1.**
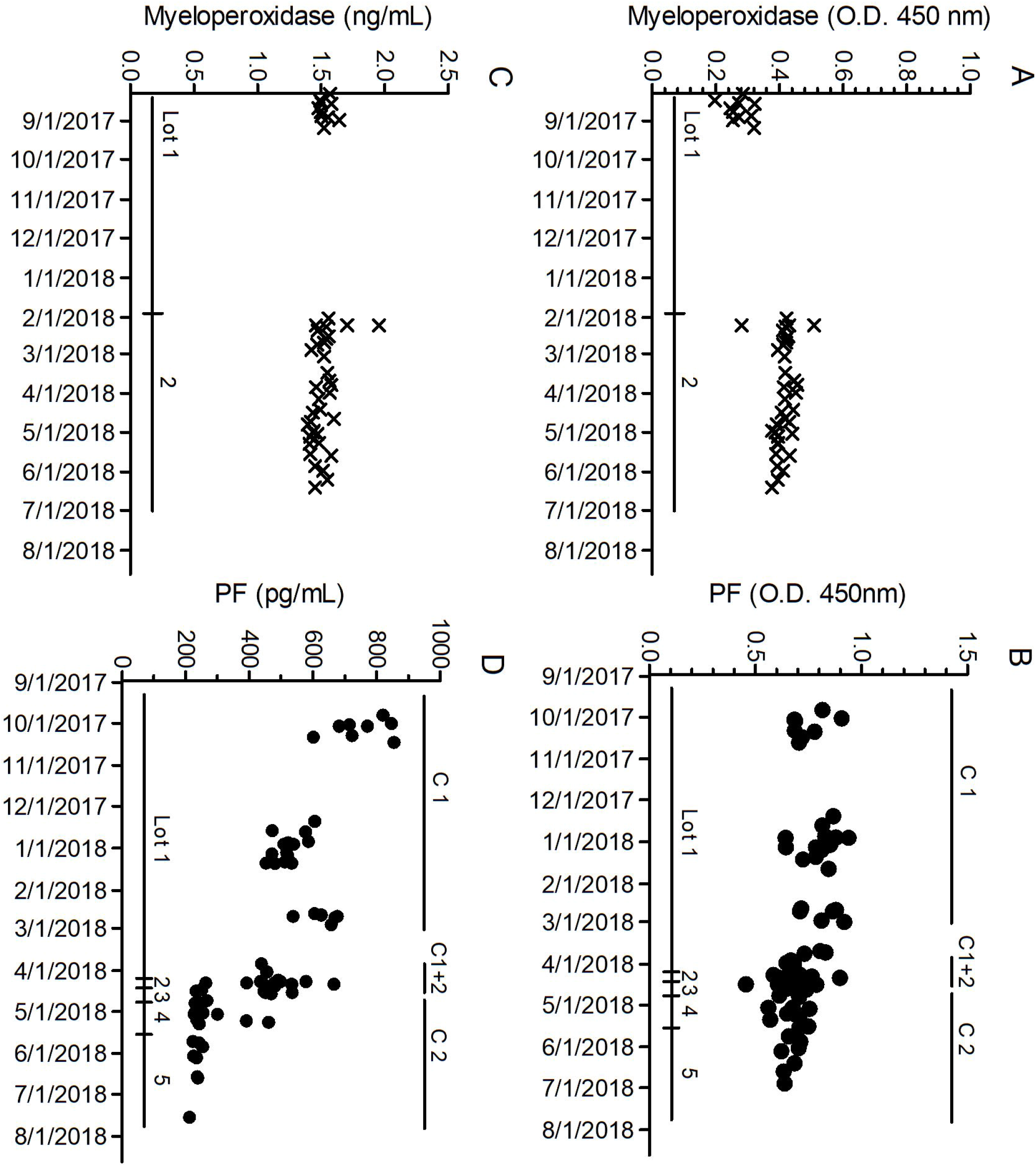
Biomarker Controls with Time. ELISAs were completed over ~10 months as described in Methods for biomarkers myeloperoxidase and PF. (A,B) Optical density (O.D. at 450nm) readings and biomarker concentrations calculated from each plate’s standard curve (C,D) for the BMC internal control samples are shown *versus* time. Two ELISA kit lots were used for myeloperoxidase, and five kit lots for PF. Two BMC control preparations were used for PF (C1 and C2), and one BMC control preparation for myeloperoxidase. OD readings for PF controls are reasonably constant with time, but unlike myeloperoxidase, PF concentrations decreased by 62.4% over the study period.

Loss of PF antigenicity in the BMC controls during freezer storage could be a contributor. BMC control preparation C1 was prepared in August 2017 and C2 in October 2017, and stored aliquots were used over the study period. The averaged optical density values for C1 were higher than C2 (P<0.01; C1 O.D.= 0.759 ± 0.095, CV=9%;n=26 plates; C2 O.D. = 0.672 ± 0.062, CV=13%; n=16 plates). However, O.D. values over time for each preparation were reasonably stable (Fig 1B), suggesting PF antigenicity did not change significantly during storage. Despite similar O.D. values, a plate with control C1 had a calculated PF concentration of 723.9 pg/mL (mean, triplicate wells) in October 2017 with kit lot #1, but the same control calculated as 238.5 pg/mL in June 2018 with kit lot #5 (Table 1). A similar change was observed for BMC control preparation C2 samples. Other than the kit itself, a detailed operations review did not identify significant changes in PF antigenicity, environment, equipment or operator contributions that could explain the large change in calculated PF concentrations observed in the BMC controls (data not shown). Notably, no similar changes were observed with the other eight biomarker kits (1-6 kit lots) over the same period.

**Table 1.** BMC Controls and ELISA Kit Lot number (#)

| Assay Date | BMC Control C1 | BMC Control C2 | Kit Lot # | O.D.<br>(450nm)* | PF Conc.<br>(pg/mL) |
| --- | --- | --- | --- | --- | --- |
| 10/11/2017 | x |  | 1 | 0.686 | 723.9 |
| 06/19/2018 | x |  | 5 | 0.638 | 238.5 |
| 02/19/2018 |  | x | 1 | 0.717 | 605.4 |
| 06/04/2018 |  | x | 5 | 0.684 | 225.1 |

\* Mean O.D. ± S.D. is $0.759 \pm 0.095$ for C1 (CV=13%, n=26) and $0.672 \pm 0.062$ (CV=9%, n=16)

In contrast, comparison of PF standard curves from kit lot #1-5 showed a trend with time (Fig 2A) that paralleled changes observed with the BMC control concentrations. Expected variability between standard curves within the lot was observed (Fig 2B), but lot-to-lot variability showed a left-shift trend. The averaged standard curves for lots #1 and #2 (September, November 2017) were similar, but lots #3-5 (April, May 2018) curves had increasing O.D. at each standard concentration. The expected consequence of a left-shift in standard curves will be lower biomarker PF concentrations, which was observed (Fig 1, Table 1).

**Fig 2.**
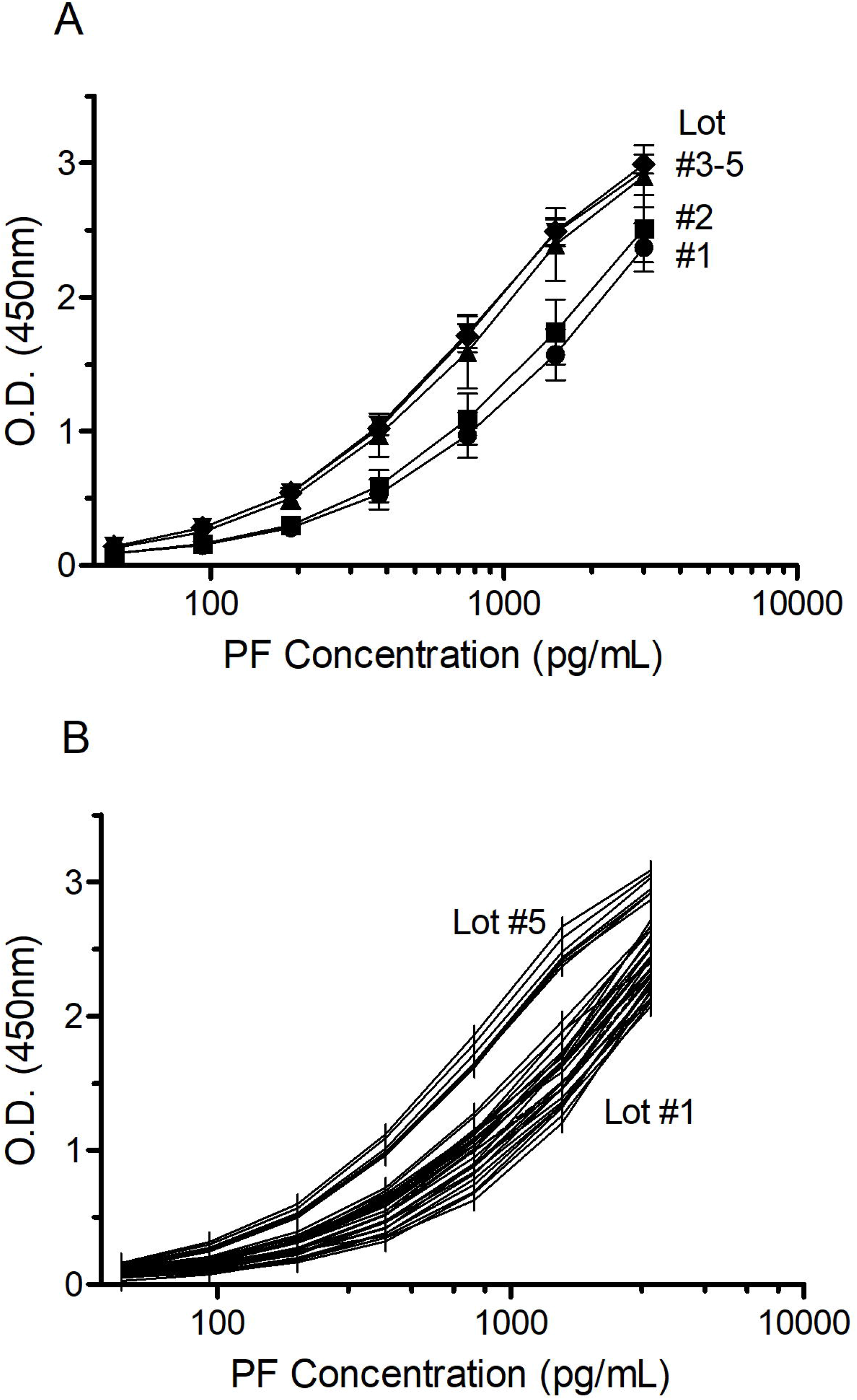
Lot-to-Lot Variability in PF ELISA. (A) The averaged standard curves for each lot of PF ELISA as the mean optical density O.D. ± S.D. at each standard concentration. N= 28, 19, 8, 4, 9 curves for lots #1-5, respectively. (B) Variability of standard curve optical density at 450nm within and between PF ELISA kit lots #1 and #5.

To address this analytic variable and rescue our patient data, a strategy was developed such that each PF standard curve is compared to a best-fit Reference curve (O.D. *versus* PF concentration) and a Shift factor (“S”) calculated that quantifies the difference between each plate’s standard curve and the Reference curve. An adjusted plate standard curve equation is derived and the biomarker concentrations are recalculated. Thus, every plate has a unique adjustment, based on the calculated S factor for that plate, and results are generated on a uniform platform. A small S factor value indicates the original standard curve on that plate is similar to the best-fit Reference curve. Conversely, a large S factor indicates a larger left-or right-shift relative to the best-fit Reference curve. The PF Reference curve was fitted with a 4-parameter logistics (4pl) equation common to many ELISA analyses, but a 5-parameter logistics (5pl) curve fit option is also available (Supplemental Methods).

Assignment of biomarker PF data to calculate the Reference curve was based on our available data, the manufacturer’s screening data, and judgment. We used data from 28 standard curves (PF kit lot #1) consistent over six months and three operators to calculate the (4pl) Reference curve. Fig 3 shows the Reference curve with unadjusted averaged standard curves for lots #2 and #5. Lot #2 curves had an average S factor of 0.0690 (n=19) indicating small differences from the Reference curve. In contrast, lot #5 curves had an average S factor of 0.6994 (n=9), indicating a substantial shift relative to the Reference curve.

**Fig 3.**
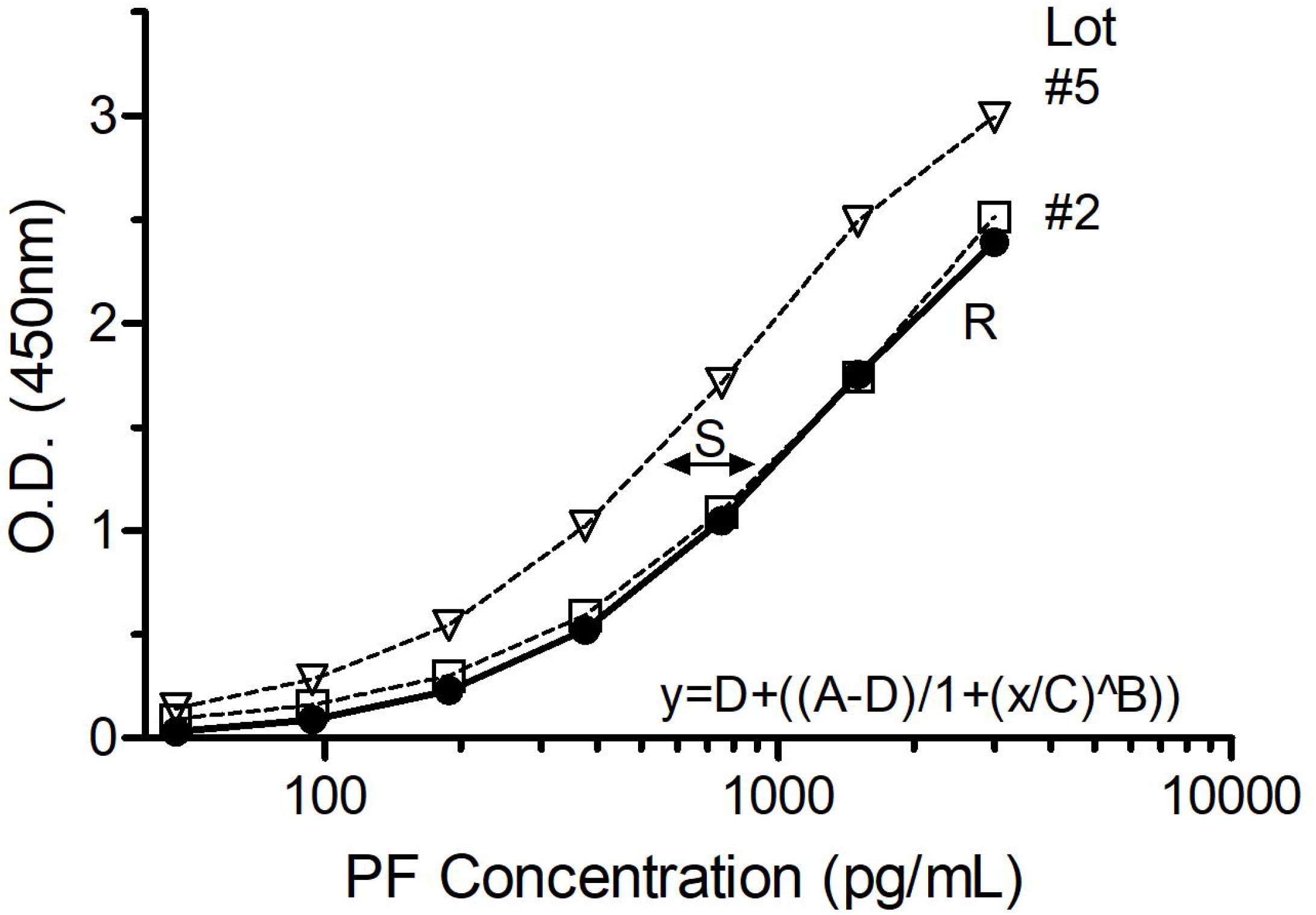
Reference Curve for calculation of S factor. The best-fit, 4-parameter logistic (4pl) Reference curve (closed circle, R, solid line) from twenty-eight PF ELISA kit lot #1 standard curves is shown as derived from the indicated equation and D= −0.01; A= 3.20; C= 1300.00; B= −1.30. The averaged standard curves for kit lot #2 has an S factor = 0.0690 (open square; n=19 curves) and S= 0.6994 (open triangle; n=9) for curves from kit lot #5.

S factors for each PF standard curve were calculated (Fig 4). Data from lot #1 was used to fit the Reference Curve, so the average S factor is close to 0 as expected. A shift from the Reference curve for lots #3-5 is shown by their higher average S factors. One lot #2 plate had an S = 0.7032, which is 20.8-fold higher than the average S factor from the remaining plates (mean S=0.0338 ± 0.158; n=18). Data from this plate is under review. Average S factors for lots #3-5 are similar, but those from lot #3 have greater inter-assay variability. The CV for lot #3 S factors is 44.2% (n=8), compared to 16.7% (n=4) for lot #4 and 11.5% (n=9) for lot #5. Whether this is due to manufacturer’s differences, sample size or laboratory-based variables is not known.

**Fig 4.**
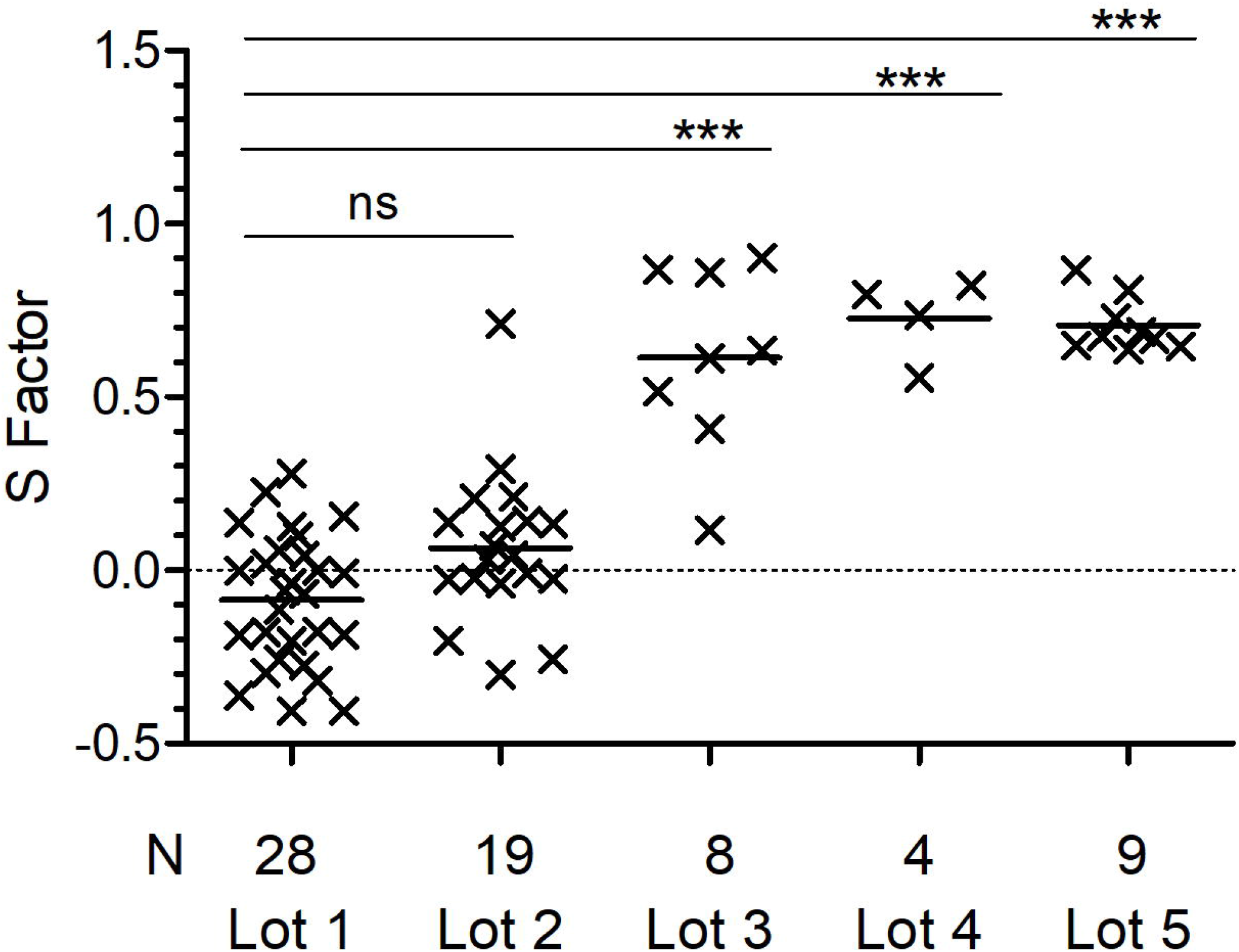
Calculated Shift Factors for PF ELISA Standard Curves. A 4pl Reference curve was derived from PF kit lot #1 standard curves data and a Shift factor “S” was calculated to quantify the difference between each plate’s standard curve and the best-fit reference curve. The S factors for each curve in lots #1 −5 is shown with the mean (horizontal line) for each lot. S factors for lots #1 and #2 curves are not different, but those for lots #3-5 differ significantly from lot #1 (***P<0.0001, ANOVA with Bonferroni post-test).

The BMC control PF concentrations for preparations C1 and C2 were re-calculated using the S factor for each plate’s standard curve. The reference curve was based on kit lot #1 data and C1 was assayed primarily with kit lots #1 and #2. Thus, the average PF concentration did not change significantly (P=0.843), but the variance was reduced (P= 0. 002), as expected (Fig 5). C2 controls were included primarily on plates from kit lots #3-5, so the difference after correction with S factors is significant (P<0.0001).

**Figure 5.**
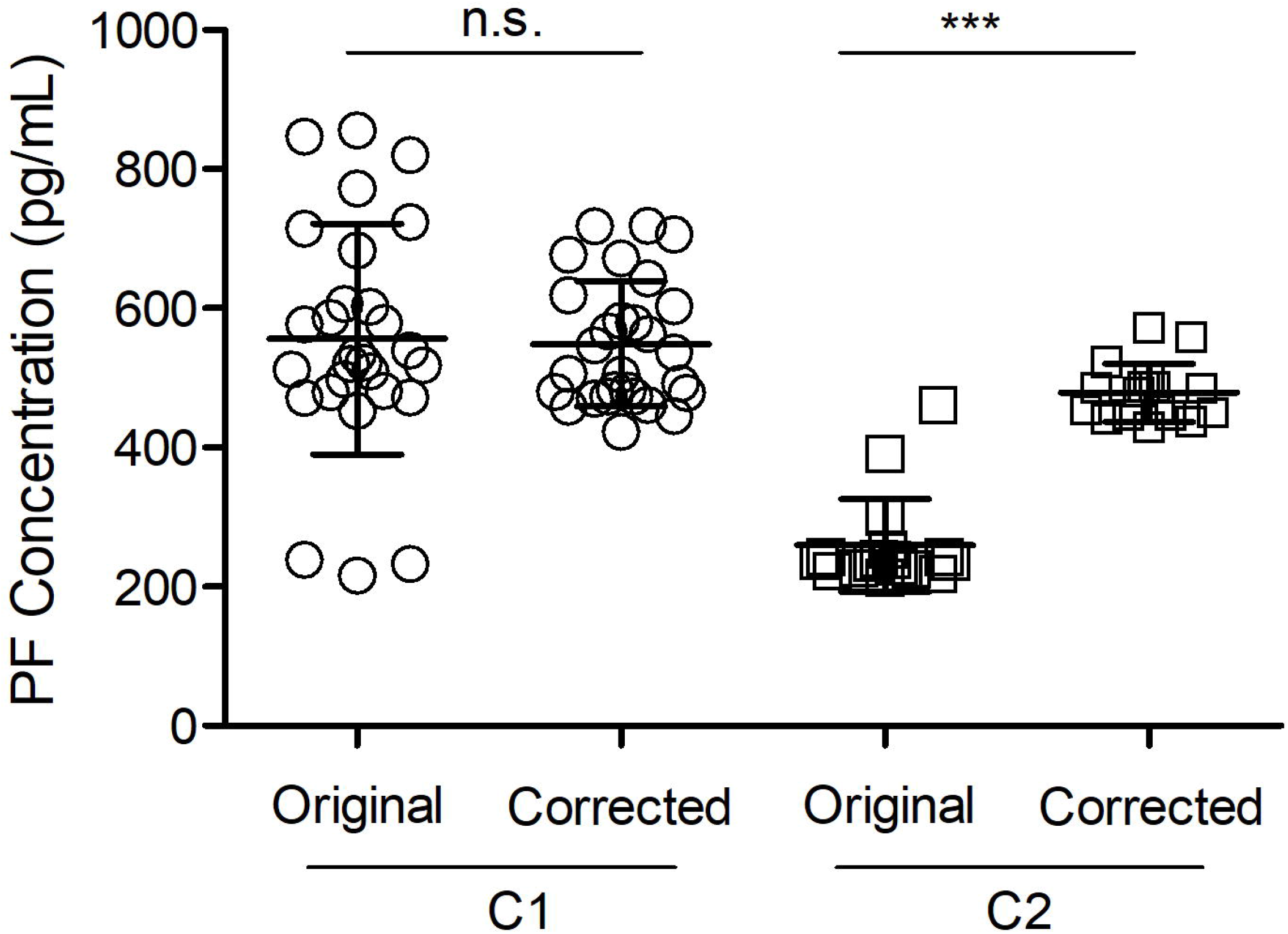
BMC Control PF Concentrations. A BMC control sample from preparation C1 or C2 was included on each ELISA plate for the 10 month study period and assayed with kit lots #1 −5. C1 was used primarily on plates from kit lots #1 and #2. C2 was used primarily on plates from kit lots #3-5. Standard curves from each plate were adjusted according to their calculated S factor and the control PF concentrations were re-calculated. The data shows the PF concentrations before and after correction with the S factors. ***P<0.0001; ns, not significant (Student’s t test).

S factors for myeloperoxidase, soluble P-selectin and plasminogen activator inhibitor-1 were calculated (Table 2). Data for 4pl Reference curves was chosen from four standard curves performed by 3 different operators over at least 6 months to represent the composite data and inter-assay variability. One myeloperoxidase curve had an S = 1.099, and this data is under review. All BMC controls were within quality control limits. Overall, their calculated S factors agree with consistent BMC Control values over time and between kit lots.

**Table 2.**
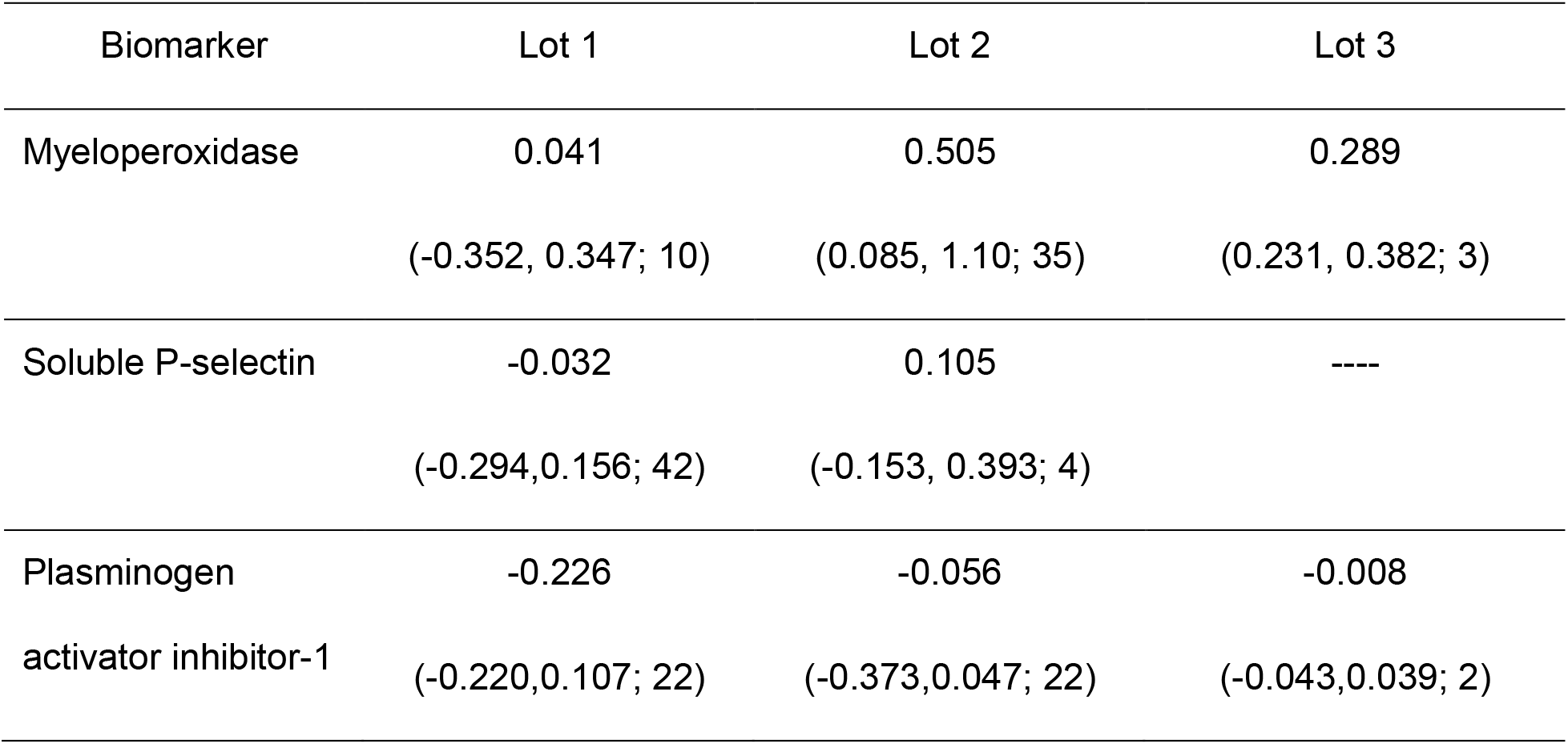
Average S Factors (min, max; N) for Biomarker Standard Curves from Kit Lots.

| Biomarker | Lot 1 | Lot 2 | Lot 3 |
| --- | --- | --- | --- |
| Myeloperoxidase | 0.041<br>(-0.352, 0.347; 10) | 0.505<br>(0.085, 1.10; 35) | 0.289<br>(0.231, 0.382; 3) |
| Soluble P-selectin | -0.032<br>(-0.294, 0.156; 42) | 0.105<br>(-0.153, 0.393; 4) | ---- |
| Plasminogen<br>activator inhibitor-1 | -0.226<br>(-0.220, 0.107; 22) | -0.056<br>(-0.373, 0.047; 22) | -0.008<br>(-0.043, 0.039; 2) |

## Discussion

Acquiring ELISA data using commercial research kits over a prolonged period presents challenges for quality control needs. There is potential impact of multiple operators, antigen stability, environmental and equipment drift, lot-to-lot reagent variability, and lack of validated controls [15, 17–20]. It is virtually impossible to manufacture a new reagent lot that is identical in all respects to prior lots, and reagent variability occurs even with commercial diagnostic assay reagents [21]. Yet minimizing these influences is necessary so patient or other experimental data can be compared with confidence over the study period. In this study, changing BMC control concentrations for biomarker PF first raised the potential of excessive lot-to-lot variability in the ELISA kits. The optical density data did not change significantly, but calculated PF concentrations greatly decreased, and this disconnect triggered the comprehensive data review. After excluding other reasonable contributors, the S factor strategy was developed to provide a uniform platform for comparison of patient PF data collected over many months.

Calculation of S factors allowed retrospective adjustment of each plate’s standard curve. The averaged S factors for PF kit lots #1-5 ranged from −0.086 to 0.735, and reduced the BMC control inter-assay variability to within our quality control limits (<9%). Review of the S factors for each standard curve was useful in that we could rapidly identify possible outliers. One curve each from PF ELISA lot #1 (high) and lot #3 (low), and one for myeloperoxidase (high) were sufficiently different and those plates were tagged for review. Unanticipated variability occurred even with SOPs, required documentation, equipment calibration and heightened operator awareness of pre-analytic variables. Controlling pre-analytic variables for quality assurance and method validation is difficult [22] and identifying trends on a day-to-day basis is largely subjective. Once a Reference curve is established, calculating the S factor for each curve provides rapid and quantitative comparative data.

Few research-use ELISA kits include validated controls, so the early decision to include spiked plasma controls on every plate was advantageous. One limitation of laboratory-made controls is that each preparation will be slightly different and those differences may not be apparent until enough plates have been run to set the quality control ranges. Access to independently standardized and validated controls in a variety of matrices (plasma, serum, urine, etc) for established and emerging biomarkers would be a valuable resource for investigators and facilitate more consistent biomarker results between laboratories. This in turn supports the goals of improved data rigor and reproducibility, and the more ethical discussion regarding disclosure of validated data to study participants [23–25].

The problem of inter-assay variability is not new [26] and various approaches are proposed to quantify data that are acquired in batches using conversion of a signal from known samples into a meaningful value for unknown samples [15, 27]. For *in vitro* diagnostic ELISA kits used in a clinical laboratories, quality assurance is provided by a validated control sample(s) with defined value limits. Research-only ELISA kits do not have this foundation, yet are used universally. Calculation of S factors and retrospective re-analysis with adjusted standard curves is useful when quality control values exceed limits and environmental, procedural, equipment or operator contributions are ruled out.

This strategy should be used judiciously primarily because data choices for the Reference curve is partially subjective. We chose data from PF ELISA lot #1 based on volume of data, relative consistency over many months and similarity to the manufacturer’s data for that lot. We could have chosen standard curve data from lot #5 plates instead, but there were fewer curves over a shorter time period and our collective experience judged this to be a less favorable choice. We do not anticipate using this strategy for the other biomarkers. Their inter-assay ELISA variability is acceptable, their BMC controls remain within limits, and there is no justifiable need to adjust the curves. That said, we have found the S factors to be a useful monitor of ELISA outcomes to rapidly identify pre-analytic factors, such as operator differences, that otherwise may be difficult to identify by visual inspection of the data, particularly for long-term projects.

There are other approaches to adjust for batch effects in immunoassays similar to our current work. More complicated statistical approaches are employed, such as the mix-effect model [15] and the iterative maximum likelihood method [27]. They assume a linear relationship between the measured signals and the analyte concentrations, which is only an approximation and has its own limitations. Our implementation takes the form of a nonlinear logistic function. Our proposed statistical model is simple and the assumption is appropriate for the observed lot-to-lot variability (Supplementary Figure 2).

We focused on developing an accessible and generalizable strategy to solve similar issues as long as the assumptions are met. The software is implemented in R and the data input uses a format familiar to those who use standard 96 well plates for ELISAs. It is written for either Mac or PC and a choice of 4pl or 5pl curve fitting is provided. The software is open source and in the public domain, with instructions for data input in the Supplemental Methods.

## Supporting information

## Acknowledgements

The authors gratefully acknowledge Dr. Thomas Kepler (BUSM, Department of Mathematics and Biostatistics, Department of Microbiology) for strategic discussions and Dr. Daniel Remick (BUSM, Department of Pathology and Laboratory Medicine) for critical reading of the manuscript. We gratefully acknowledge the administrative assistance of Lindy Joseph (BUSM) and Dr. Jasmin Bavarva (Leidos Biomedical Research, Inc., Frederick, MD) who contributed to project set-up.

## Supporting Information

Supplemental Methods provides instructions on how to load and use ELISAtools for calculation of S factor(s).

