## Supplementary material for "A computational solution to improve biomarker reproducibility during long-term projects"

**Supplemental Methods**

Summary: To calculate S factors, the user will engage ELISAtools and this program uses R code provided in “tutorial”. Four input .txt files need to be provided by the user. A simple way to do this is to put the data in Excel and save as .txt files. Example files are given in the ELISAtools package. The following is a guide for the steps necessary to obtain S factors. There are some differences between PC and Mac users, and note that R uses unformatted text and is very specific about format. A 5pl curve fit is the default. To change to 4pl, see instructions in Troubleshooting Tips at the end.

The ELISAtools package is available from: <https://github.com/BULQI/ELISAtools>

**Part I. Creating the Input Files**

The data for the Reference standard curve must be chosen by the user. In practice, we use data from at least 4-6 standard curves, preferably from several ELISA kit lots (aka, batches) and multiple operators to represent expected variability.

Terminology: *kit lot* refers to the commercial kit lot number assigned by the manufacturer. *Batch* is the ELISAtools equivalent of a kit lot. *Batch-corrected* is the final adjusted concentrations based on that plate’s standard curve adjusted with its S factor.

Four input files will be read during the analysis. These are the Design, Annotation, Standard Concentration and Optical Density files. It is easier to keep these file in one single folder, such as the default data folder “ELISAtools/inst/extdata”. With the help of the Design file, the Annotation and Standard Concentration files direct R to assign identities for each well to correspond with O.D. readings in the wells of the Optical Density file. Correct syntax and formatting within these files is important.

1. Design File. The design file pulls information from the annotation, standard concentration and optical density files. There is one design file per ELISAtools analysis.
   1.1 In the ELISAtools folder (on your desktop), open the design.txt file. [ELISAtools>inst>extdata>design].
   1.2 The easiest way to get started is to copy/paste this default design text into an Excel worksheet. In Excel, the first row is the ‘header’. Do NOT change this first line of text otherwise the program will not run. The file is read line by line for details of a particular assay, such as the experiment ID, name of raw data file, date of assay, standard concentration.

In Excel, for rows 2-4 (or more) and columns A-I, you can change the text to customize to your particular set of experiments or data.

Column B (FileName) shows the names of the raw OD files.

Column C (Batch) is the lot numbers/names.

Column D (Num_Plate) shows the number of ELISA plate data in each file

Column E (Date) is the date!

Column F (AnnotationFile) shows the name of the annotation file you will make

Column G (Std_Conc) is the file(s) you will make with your standard concentrations

Columns H and I do not have to be used. These are for reading files that are not saved in the default “extdata” folder.

1.3 Name the Excel file (e.g., mydesignfile.txt) and save it as a .txt file in the extdata folder within ELISAtools [ELISAtools>inst>extdata].
1.4 IMPORTANT: The first experiment entered in the design file is automatically designated your Reference curve. If multiple curves are desired to create the Reference curve, you have to designate additional experiments *with the same batch name*. Column C of the design file is “batch”, so copy the name of the batch (e.g., BatchRef) into each field of that column as desired, and those data will be used for the Reference curve.
    1.4.1 The code “ref.batch=1” is default within the program, but can be changed if another batch should be used as reference.
1.5 We recommend to save the Excel file in the extdata folder as a workbook in the Excel format because you can easily edit it. You can re-export the .txt files into the extdata folder as needed. Be sure to use: File>SaveAs>Excel workbook. It helps to rename the tabs for the files (myDesignfile, Annotation, Std Conc).

2. Annotation file. This file tells the program what is in each well. Again, an example is provided in ELISAtools and is easy to download for user modification. The format should not be changed. You can have an annotation file for each plate, or for multiple plates if they share well designations and layouts.
   2.1 Open the AnExp_2plate.txt file (in the extdata folder). Copy/paste the text in the Excel file on a different worksheet. You will see a familiar ELISA plate map. You can edit the well names to reflect your experiment.
   2.2 The standards are noted as s1-s8. You can move these anywhere you wish, but do NOT change the name. They must remain s1-s8 format. If you have five standards, then use s1-s5, etc.
   2.3 Save the annotation file in the extdata folder (e.g., my_annotationfile.txt). If you have (say) five plates, copy/paste the format vertically and re-name each plate (plate 3, plate 4, plate 5). Be sure to have “~End” to mark the end of each plate.
3. Standard concentrations file. This provides the concentrations for each standard curve that is used for the particular ELISA. You can have a standard concentration file for each plate, or for multiple plates if they share the same standard concentrations.

3.1 Open the stdConc.txt file (in the extdata folder). Copy/paste the text in the Excel file on a different worksheet.
3.2 Change the standard concentrations if you wish for each plate. Copy/paste vertically for multiple plates and change the plate name accordingly.

4. Optical Density File.
   4.1 Now input your raw data. You’ve run your ELISAs and you’ve exported the optical density data into a .txt file(s). Save the file(s) directly in the extdata folder within ELISAtools. The names of these files should match the names in Column B of your design file (mydesignfile in our example).
5. Final Check: In the Excel workbook, go to the first myDesignfile tab. Confirm that all the names are correct in column B and that they end in .txt. Repeat the review for annotation files (column F) and standard concentration files (column G). Update and re-export this mydesignfile.txt (in the extdata folder).

**Part II. Customizing the Code**

1. R is an open source software. Download version 3.5.1 or later onto your computer and open it.
2. Download the ELISAtools package folder from https://github.com/BULQI/ELISAtools to your desktop
   2.1 Rename the “ELISAtools-Master” folder to “ELISAtools” (remove “-Master”). Review the files in the package. Alternatively, you can download and install the package directly by running in R the command "devtools::install_github(‘bulqi/ELISAtools’)".
   2.2 Open the tutorial file within R using the open folder icon or: [Desktop>ELISAtools>inst>extdata>tutorial].
   2.3 Two windows are open: R Console and R Editor. R Editor currently displays customized code (known as tutorial) for ELISAtools.
3. User-specific modifications to the tutorial code need to be made the first time ELISAtools is used. These changes are specific to your files and file locations, and the name of one user-specific input file (known as the Design file):
   3.1 With a PC:
       3.1.1 Before the top line of code, add two lines:
a) setwd("*put your folder location here*").

As an example: setwd("C:\\Users\\dsk\\Desktop")

b) install.packages(c("ELISAtools"),repos=NULL, type="source")

Skip this step if you have called "devtools::install_github" in the step 2.1

3.1.2 Technical tip: R uses unformatted text, so be sure your quotes are unformatted (use this: "). If Word insists on formatting, you will need to manually type in the command with unformatted quotes.
3.1.3 After “library(ELISAtools)”, add in:

a) system.file("extdata", package="ELISAtools")
   3.1.4 You will change 2 lines of code. The current directory commands are:

dir_file<-system.file("extdata", package="ELISAtools")

#dir_file<-("E:\\feng\\LAB\\hg\\ELISA\\ELISAtools\\dev\\morganFiles")

[Tip: The “#” symbol, when placed before a command or code, tells the program to ignore the text after the “#” within the same line.]

Place a “#” before the first directory command and remove the “#” from the second directory command to designate which command R should follow.

Thus, the second directory command will change to:

a) dir_file<-("*put your location here for your extdata directory using \\ as needed*")

Example: ("C:\\Users\\dsk\\Desktop\\ELISAtools\\inst\\extdata")

3.2 With a Macbook: Running the code with a Macbook will require minor deviations in syntax from the above instructions.
    3.2.1 For example, when specifying the directory, use forward slash “/” rather than backslashes “\\”; e.g. “/Users/macbook/Desktop”
    3.2.2 To easily copy the path file, right click on the folder and select “Get info”.
    3.2.3 Copy the path file the path file after “Where:”
3.3 With PC or Mac, you should change the name of the Design File input.
    3.3.1 Modify the name *design.txt* within “batches<-loadData(file.path(dir_file, "design.txt"))" to reflect the name of your design file. Remember to include the file extension as .txt
    3.3.2 If you leave it with the name *design.txt*, it will run the default design file.
3.4 Save your changes: Click on the R Editor window, then the Save Script icon on the ribbon.

4. In R editor: When you’re ready to run your customized code, accessory packages need to be installed from R servers to properly run ELISAtools:
   4.1 With a PC:
       4.1.1 In the R Console before running the code, at the cursor add in:

utils:::menuInstallPkgs()

4.1.2 Press Enter if needed
4.1.3 R Console will prompt you to select a CRAN mirror for use in this session from a pop-up window.
4.1.4 Select a CRAN mirror – Select one that is geographically close to you. Hit OK if needed. For this example, one of the USA mirrors was used.
4.1.5 Select “R2HTML”from the menu
4.1.6 Click OK
4.1.7 Answer Yes to personal library (if it comes up). Answer Yes to create destination of the personal library.
4.1.8 Use the utils:::menuInstallPkgs() command two more times to select for “minpack.lm”, and “stringi”
4.2 With a Macbook:
    4.2.1 Select Packages & Data from the top of the program.
    4.2.2 R Console will prompt you to select a CRAN mirror for use in this session from a pop-up window.
    4.2.3 Select a CRAN mirror – Select one that is geographically close to you. Hit OK. For this example, one of the USA mirrors was used.
    4.2.4 Search for the following packages and then hit install: “R2HTML”,“minpack.lm”, and “stringi”

5. Go to the R Editor window. Set your cursor at the very beginning, select all the code and copy/paste to the red cursor in the R Console window.
6. Now the program starts to read. You may have to press Enter before it starts to read. You will see a series of graphs as it works and generates curves and reports. You know the program is finished when you see the blinking cursor at the bottom. You may need to press Enter to run the last line of code.
7. When finished you will see:

*** Output redirected to directory: .

*** Use HTMLStop() to end redirection.

**Part III. Getting the Reports**

1. Go to the folder you specified in the dir_file<-("put your location here for your extdata directory using \\ as needed"). Open the report created: report_com_ana.html (there is also a .txt version). This is the combined report from the individual batch reports.
2. It is helpful to rename this file before you run the program again, otherwise it will re-write over your current report. (You could also move all the report files to a different folder). Double click on it and it will open in a web browser.
3. You will see the ELISA Data Analysis Tool Report. The beginning of the report has the Model Parameters (equation with parameters), S Factors for each batch, and a graph of all the adjusted curves together.
4. Individual reports for each batch are next. For each there is a graph of the raw data curves with the fitted batch mean. A table includes the ID, plate locations (row, column), raw OD, the original concentration (conc_pred) and the adjusted concentration based on the best-fit Reference curve (conc_pred.bc).
5. If you opened the report as a text file, you will see basically the same thing but without the graphics. It is possible to open this file in excel, and from there, sort or filter data by batch, control ID, or date.

******************************************************************************

Troubleshooting Tips:

- R analysis requires very strict input. An error such as “plate not found” or ‘Batches does not exist’, could mean there is an issue with the input files or the location of the files.
  - Adding all relevant files to the ‘extdata’ folder of the package should solve this issue.
  - Alternatively, if input files are developed on a Macbook, it may be helpful to confirm that the input is in .txt and not Unicode variations.
  - Look for additional or incorrect characters within the files.
  - Make sure you have the permission to write output files to your working folder.
- Several options can be found within the code and are ignored by the “#” symbol. For example, both 4pL and 5pL options are available, but since 5pL is the default, the code for 4pL analysis is blocked by using a # before these commands. If a 4pL curve fit is desired, the user can place the “#” before 5pL code and delete them from before the 4pL code. Here is an example of where the code should be modified to run 4pL code:

model<-"5pl"

pars<-c(7.2,0.5, 0.015) #5pl inits

names(pars)<-c("xmid", "scal", "g")

#model<-"4pl"

#pars<-c(7.2,0.9) #4pl inits

#names(pars)<-c("xmid", "scal")

batches<-runFit(pars=pars, batches=batches, refBatch.ID=1, model=model )

#batches<-runFit(pars=pars, batches=batches, refBatch.ID=1, model="4pl" )

- Change to:

#model<-"5pl"

#pars<-c(7.2,0.5, 0.015) #5pl inits

#names(pars)<-c("xmid", "scal", "g")

model<-"4pl"

pars<-c(7.2,0.9) #4pl inits

names(pars)<-c("xmid", "scal")

#batches<-runFit(pars=pars, batches=batches, refBatch.ID=1, model=model )

batches<-runFit(pars=pars, batches=batches, refBatch.ID=1, model="4pl" )

~End
